# Complementary information on single nucleotide variants, INDELs and functional translocations can be obtained with RNAseq using different library preparations

**DOI:** 10.1101/301010

**Authors:** Riccardo Panero, Maddalena Arigoni, Martina Olivero, Francesca Cordero, Alessandro Weisz, Marco Beccuti, Mariaflavia Di Renzo, Raffaele A. Calogero

## Abstract

**Background:** RNA-seq represents an attractive methodology for the detection of functional genomic variants because it allows the integration of variant frequency and their expression. However, although specific statistic frameworks have been designed to detect SNVs/INDELS/gene fusions in RNA-seq data, very little has been done to understand the effect of library preparation protocols on transcript variant detection in RNA-seq data.

**Results:** Here, we compared RNA-seq results obtained on short reads sequencing platform with two protocols: one based on polyA^+^ RNA selection protocol (POLYA) and the other based on exonic regions capturing protocol (ACCESS). Our data indicate that ACCESS detects 10% more coding SNV/INDELs with respect to POLYA, making this protocol more suitable for this goal. Furthermore, ACCESS requires less reads for coding SNV detection with respect to POLYA. On the other hand, if the analysis aims at identifying SNV/INDELs also in the 5’and 3’ UTRs, POLYA is definitively the preferred method. No particular advantage comes from the usage of ACCESS or POLYA in the detection of fusion transcripts.

**Conclusion:** Data show that a careful selection of the “wet” protocol adds specific features that cannot be obtained with bioinformatics alone.

## Background

Whole Exons Sequencing (WES) is the preferred method to detect Single Nucleotide Variants (SNVs) and intermediate insertions/deletions (INDELs) in the DNA of pathological samples. On the other hand, RNA sequencing (RNA-seq) is the method of election for gene/transcript quantification, which has nearly completely replaced expression microarrays.

RNA-seq is instrumental to detect functionally important SNV/INDELs, such as actionable mutations. INDELs represent another interesting type of variants also detectable by RNA-seq [1]. Obviously, it is also the only alternative when WES is not feasible. Unfortunately RNA-seq shows computational criticalities in SNV/INDELs detection, such as those due to splicing [2] and the need of statistical models that are insensitive to variability in read coverage due to unequal transcript expression levels [3–5]. RNA-seq has been also used for the detection of translocations generating functional aberrant proteins (also known as chimeras or fusion transcripts [6]), which could act as driver mutations in cancer [7, 8]. Indeed, the sequencing coverage required for fusion transcripts detection in RNA-seq is much lower than the one needed for Whole Genome Sequencing (WGS) making the RNA-seq a suitable methodology for fusion detection. Furthermore, many computational methods are available for the detection of fusion transcripts [9–11] using RNA-seq.

As mentioned above, most of the RNA-seq issues have been addressed using bioinformatics approaches. However, bioinformatics is not the only variable involved in variants identification in RNA-seq, since RNA-seq data can be generated with a plethora of different library preparation protocols, e.g. stranded/unstranded, rRNA depletion/polyA^+^ transcripts selection/CDS capturing, etc. To understand the effect of different library preparation protocols on variants detection in RNA-seq data, here we compare two methods: an unstranded polyA^+^ selection protocol (from now on called POLYA) and a stranded exon-specific capture protocol (from now on called ACCESS). POLYA was the first protocol designed to quantify poly-adenylated mRNAs [12], through the use of oligo-dT selection of polyA^+^ transcripts and subsequent sequencing via chemical fragmentation and random examers mediated retrotranscription; thus it does not provide strand information. ACCESS has been designed to work with “difficult samples”, characterized by RNA degradation (e.g. Formalin-Fixed, Paraffin-Embedded tissue samples [13]). It implements the oligonucleotides capturing technology used for selective selection of exons by Illumina for WES analysis, thus allowing a uniform capture of coding transcriptome and providing strand information as well. Our aim is to understand which of the two protocols provides the most robust approach to call SNV/INDELs and fusion transcripts by means of RNA-seq.

## Methods

### Data set preparation

#### Illumina MCF7 data

POLYA and ACCESS RNA-seq data derived from the breast cancer cell line MCF7 were kindly provided by Drs G. Schorth and S. Gross from Illumina (Illumina, San Diego, CA), Briefly, RNA-seq data were generated from MCF7 total RNA acquired from BioChain (BioChain Institute, Inc. Newark, CA, USA), RNA libraries were prepared according to manufacturer’s instructions using TruSeq RNA Sample Prep Kit v2 and TruSeq RNA Access Library Prep Kit (Illumina, San Diego, CA) sequenced on a NextSeq 500 sequencer in 75-bp paired end sequencing mode following manufacturer instruction. For each library preparation two NextSeq flow cells were used. We refer to the above datasets as POLYA-I and ACCESS-I. ACCESS-I and POLYA-I were combined to generate 5 data samples (s001, s012, s123, s1234, sAB1234), where the number of reads progressively increases from s001 to sAB1234 (Supplementary Table 1).

#### Mayo Clinic MCF7 WES data

MCF7 exome data set was kindly provided by Dr Y. Asmann (Mayo Clinic, Jacksonville, Florida). The data set was part of the paper published by Wang and coworkers in Bioinformatics in 2014 [1]. Exome sequencing was generated on the exome DNA fragments captured using the Agilent’s SureCapture kit v2, and sequenced on HiSeq 2000 in 100-bp paired end sequencing mode following manufacturer instruction. We refer to the above dataset as EXOME-M. The sequencing depth and mapping statistics of the EXOME-M are summarized in Supplementary Table 1.

### Data preprocessing

Data preprocessing for SNVs and INDELs detection was done essentially as described by GATK best practice (Fig. 1), no specific preprocessing was done for transcripts expression quantification and fusion detection.

**Figure 1:**
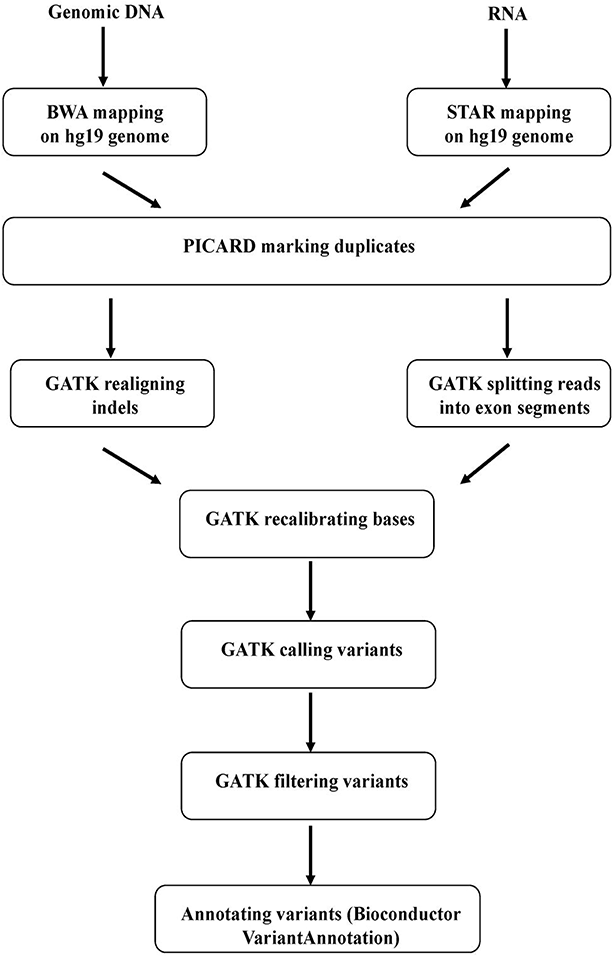
Workflow for WES and RNA-seq analysis of SNV/INDELs.

#### Exome analysis

In brief, WES fastq files for EXOME-M were mapped to human genome assembly hg19 with BWA (version 0.7.12), duplicated reads were marked with PICARD (version 1.133), GATK (version 3.5.0) was used to realign INDELs and recalibrate bases. MCF7 SNVs were detected using GenomeAnalysisTK implemented in GATK java suite. Subsequently SNVs were filtered using the following parameters, as suggested by GATK (http://gatkforums.broadinstitute.org/gatk/discussion/2806/howto-apply-hard-filters-to-a-call-set):

- QD ≥ 2, where QD indicates variant confidence divided by the unfiltered depth of non-reference samples.
- FS ≤ 60, where FS indicates Phred-scaled p-value using Fisher’s Exact Test to detect strand bias in the reads.
- MQ ≥ 40, where MQ indicates the root mean square of the mapping quality of the reads across all samples.
- MQRankSum ≥ −12.5, where MQRankSum indicates the u-based z-approximation from the Mann-Whitney Rank Sum Test for mapping qualities. This test is only applied to heterozygous calls.
- ReadPosRankSum ≥ −8, where ReadPosRankSum is the u-based z-approximation from the Rank Sum Test for site position within reads.
- QUAL ≥ 100, where QUAL is the Phred-scaled probability that a reference/alternative polymorphism exists at this site given sequencing data.
- DP ≥ 10, where DP indicates the number of filtered reads that support each of the reported alleles.

The detection of INDELs was done using the same procedure described above for SNVs detection. Furthermore, INDELs were filtered using the following parameters, as suggested by GATK (http://gatkforums.broadinstitute.org/dsde/discussion/2806/howto-apply-hard-filters-to-a-call-set):

- QD ≥ 2;
- FS ≤ 200;
- ReadPosRankSum ≤ −20

#### RNA-seq analysis

RNA-seq fastq files for ACCESS-I and POLYA-I sets were mapped to human genome (hg19) reference with two-steps STAR (version 2.3.1n), duplicated reads were marked with PICARD and GATK was used to split reads into exon segments, realign INDELs and recalibrate bases (Supplementary Table I). MCF7 SNVs were detected using GenomeAnalysisTK implemented in GATK java suite. Subsequently SNVs were filtered as described above for WES data. SNVs were annotated using VariantAnnotation Bioconductor package (version 1.16.4) [14].

The detection of INDELs was done following the recommendations described by Sun et al. [1], i.e. using the same procedure described above for SNVs detection. Furthermore, INDELs were filtered using the following parameters, as suggested by GATK (http://gatkforums.broadinstitute.org/dsde/discussion/2806/howto-apply-hard-filters-to-a-call-set):

- QD ≥ 2;
- FS ≤ 200;
- ReadPosRankSum ≤ −20

Gene/transcript expression quantification was done using RSEM (version 1.2.29) [15], hg19 genome assembly and UCSC annotation. Protocol specific expression was detected using RankProd Bioconductor package [16].

Fusion transcripts were detected using JAFFA-assembly using default parameters [17].

## Results

### Generation of datasets Error! Bookmark not defined

POLYA-I and ACCESS-I RNA-Seq data were generated from MCF7 breast cancer cells as described in the Methods section. The two short read library preparation protocols differ in the selection of RNA fragments. TruSeq RNA Library Preparation (POLYA), allows the sequencing of the fraction of RNAs characterized by a polyA^+^ tail while the TruSeq RNA Access Library Preparation (ACCESS), is based on an exons-specific capture protocol. The latter has been devised for RNA quantification in degraded samples, where polyA^+^ selection would not guarantee a good representation of full-length transcripts. The capturing technology, implemented in ACCESS, is identical to the one used for selective selection of exons in Illumina WES library preparation kits, e.g. Nextera Rapid Capture Exome protocol. Since we are interested to understand if there are protocol-specific advantages in variants detection, we used MCF7 RNA-seq data to create a set of data samples made of increasing number of sequenced reads obtained with either POLYA or ACCESS protocols (Supplementary Table 1). From both POLYA-I and ACCESS-I datasets we generated 5 sets of data for each protocol (Fig. 2 and Supplementary Table 1), made by progressive increase of reads number after duplicates removal.

**Figure 2:**
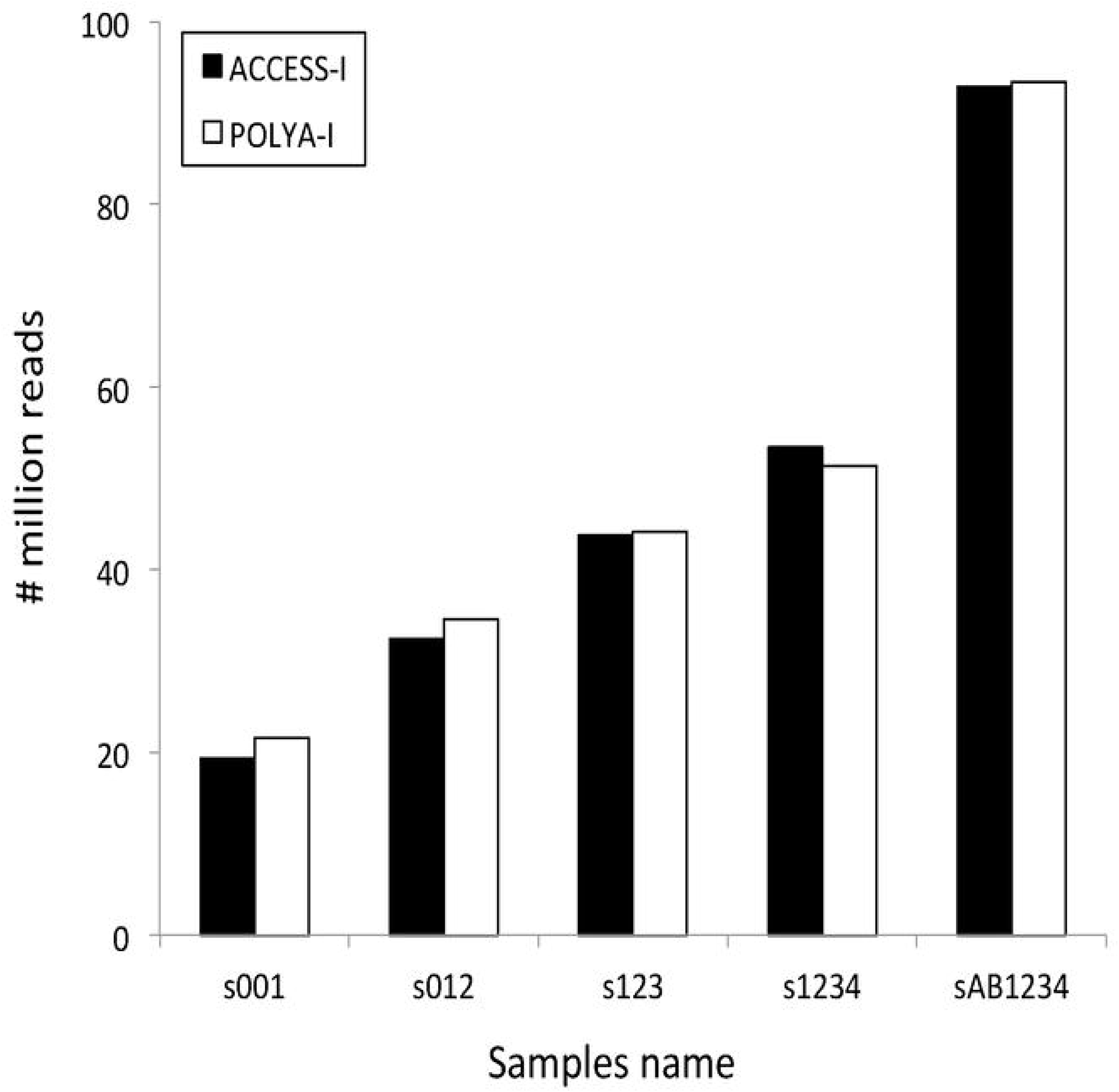
The histogram shows the mapped reads, after duplicates removal, for each of the 5 sets of MCF7 RNA-seq data obtained with either POLYA or ACCESS protocol, assembled as shown in Supplementary Table 1.

### ACCESS and POLYA provide comparable gene-level expression quantification

We first checked if any difference exists at gene-level between expression quantification done using either ACCESS or POLYA protocol. We computed gene expression using RSEM [15] and we calculated the log_2_ ratio between ACCESS-I/POLYA-I for all 5 sets. Approximately 70% of log2 ratios between ACCESS-I and POLYA-I (Supplementary Figure 1A) were within +/− 0.5 range, which represents the area of random noise for technically replicated experiments [18]. Furthermore, we evaluated the presence of statistically consistent expression differences between the two protocols analyzing the ACCESS-I versus POLYA-I expression ratio for each set of data using Rank Product [16].

The statistical analysis indicated that out of the 20017 expressed genes detected with both protocols, there are 1222 genes consistently more expressed in POLYA-I (Rank Product P-value ≤ 0.1) with respect to ACCESS and 1390 consistently more expressed in ACCESS (Rank Product P-value ≤ 0.1) with respect to POLYA. Moreover, only the 1222 genes consistently more expressed in the POLYA-I dataset were found expressed at least two fold higher than in the ACCESS-I dataset (Supplementary Figure 1B).

### POLYA detects more SNVs/INDELS than ACCESS in the whole mRNA, while ACCESS detects more SNVs/INDELs in the coding exons

The data samples s001, s012, s123, s1234, sAB1234 showed in Fig. 2, were used to detect SNVs following the GATK workflow for RNA-seq data (Fig. 1), SNVs were spread over promoters (Fig. 3A), intergenic regions (Fig. 3B), introns (Fig. 3C), coding regions (Fig. 3D), 5’ UTRs (Fig. 3E) and 3’ UTRs (Fig. 3F), The distribution of the SNVs with respect to the number of mapped reads shows that both ACCESS-I and POLYA-I are near reaching a plateau at approximately 100 million reads. POLYA (○) detected a higher number of SNVs with respect to ACCESS (●) for all annotation groups, but for the coding regions (Fig. 3D), SNVs detected with ACCESS in the coding regions were consistently 10% more than those detected in POLYA unless for the set containing the highest number of sequencing reads, i.e sAB1234 (Fig. 3D), where the differences in number between the detected SNVs drops to 0%.

**Figure 3:**
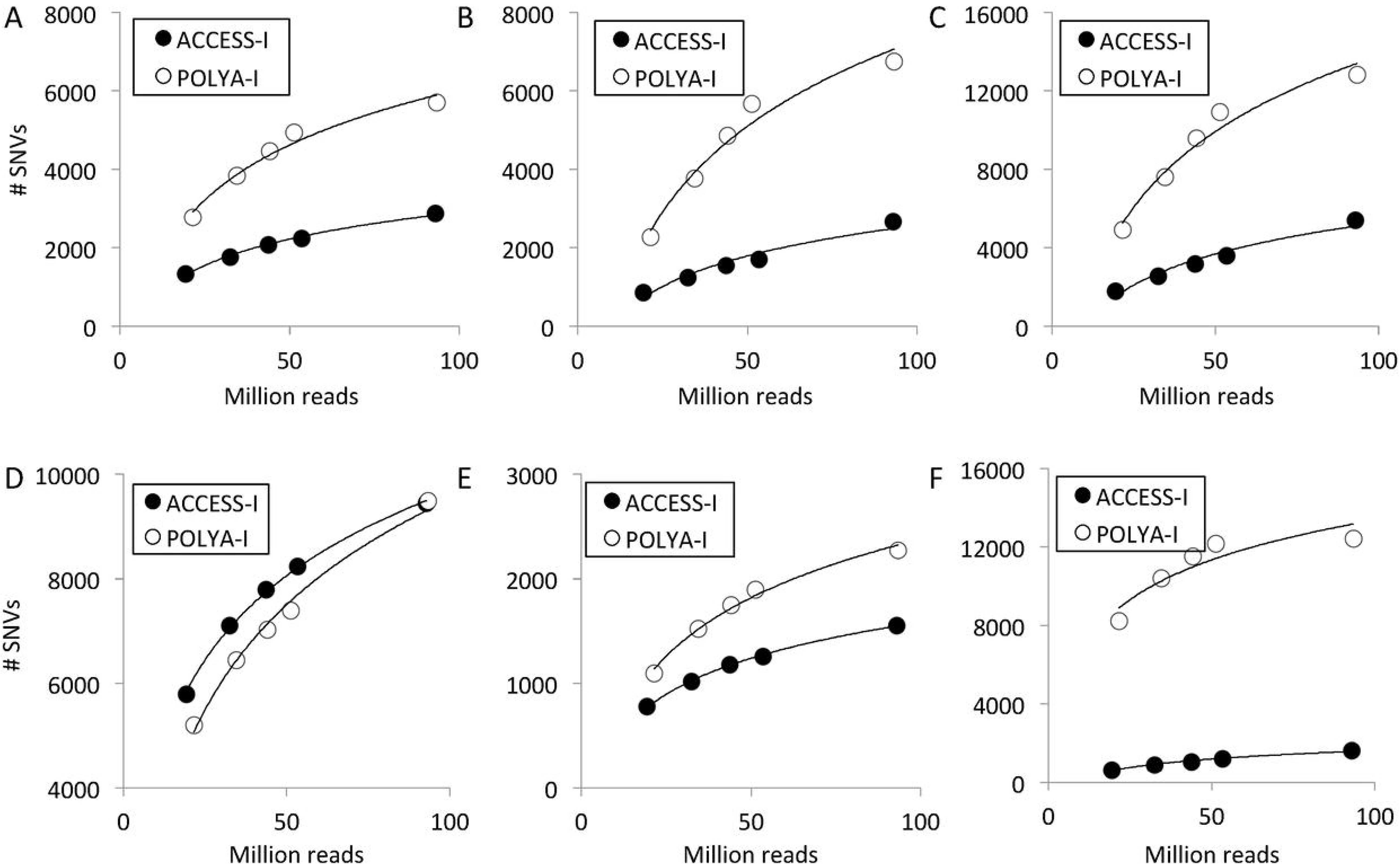
Number of SNVs detected from ACCESS-I and POLYA-I, organized on the basis of SNV location: A) Promoters, B) Intergenic, C) Introns, D) Coding, E) 5’ UTRs, F) 3’ UTRs. ○ and ● indicate, from left to right in each panel, s001, s012, s123, s1234, sAB1234 data samples.

Using the GATK workflow we also detected INDELs in the s001, s012, s123, s1234, sAB1234 sets of data samples. As shown in Fig. 4, the overall distribution of INDELs is similar to that of the SNVs. Specifically only the INDELs detected with ACCESS in the coding regions were consistently at least 25% more than those detected with POLYA unless for dataset containing the highest number of sequencing reads, sAB1234, where INDELs detected with ACCESS were only 11% more in number than those detected with POLYA (Fig. 4D).

**Figure 4:**
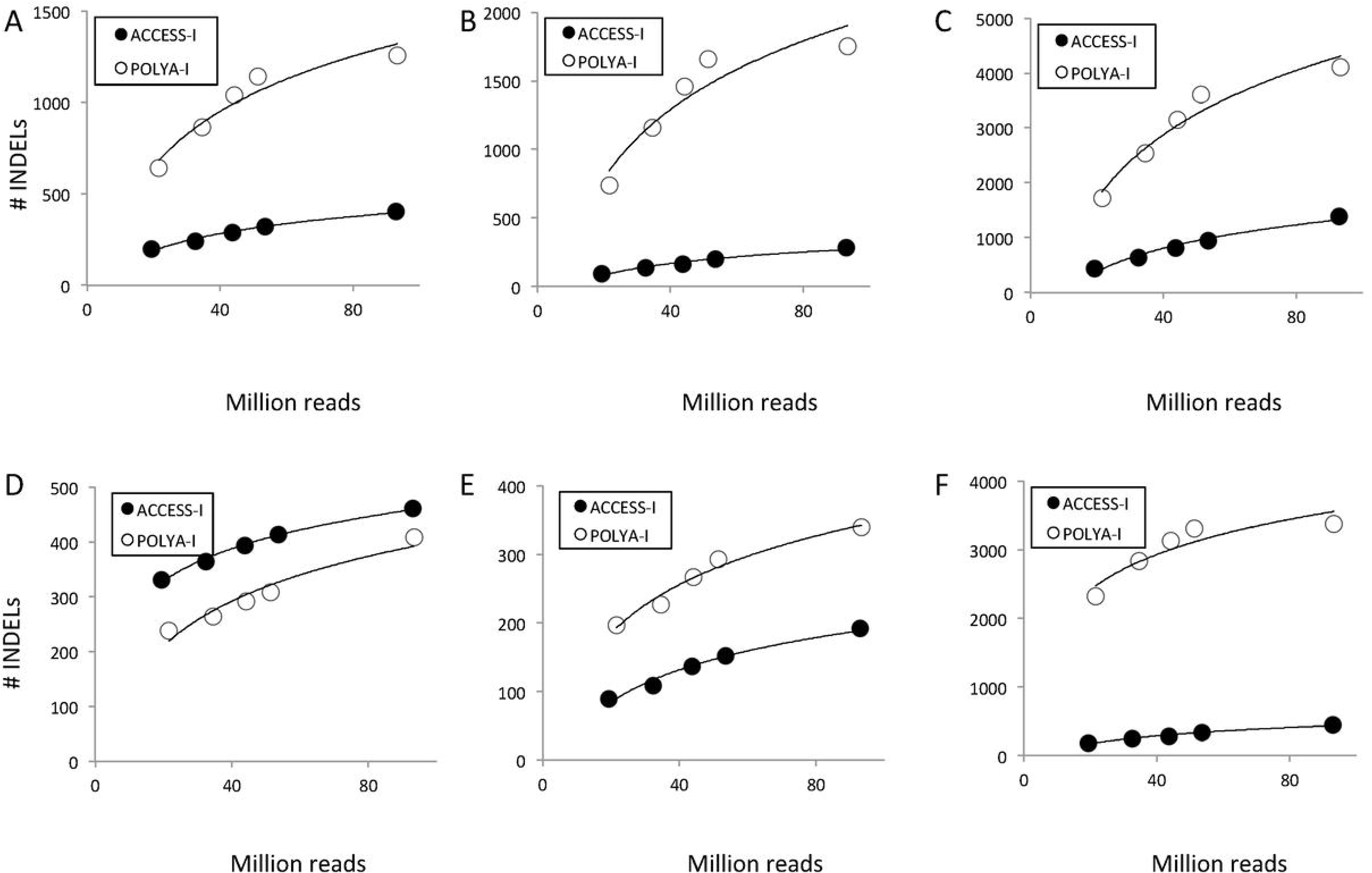
Number of INDELs detected from ACCESS-I and POLYA-I, organized on the basis of INDELs location: A) Promoters, B) Intergenic, C) Introns, D) Coding, E) 5’ UTRs, F) 3’ UTRs. ○ and ● Indicate, from left to right in each panel, s001, s012, s123, s1234, sAB1234 samples.

### ACCESS detects more coding SNVs than POLYA at low input reads

The number of SNVs, detected in common by the two protocols in coding regions, increased linearly with the increase of the number of sequenced reads (Table 1 “COMMON”, Fig. 5A □). We observed that there are approximately 10% more coding SNVs detected only by ACCESS with respect to those only specific of POLYA (Table 1).

**Figure 5:**
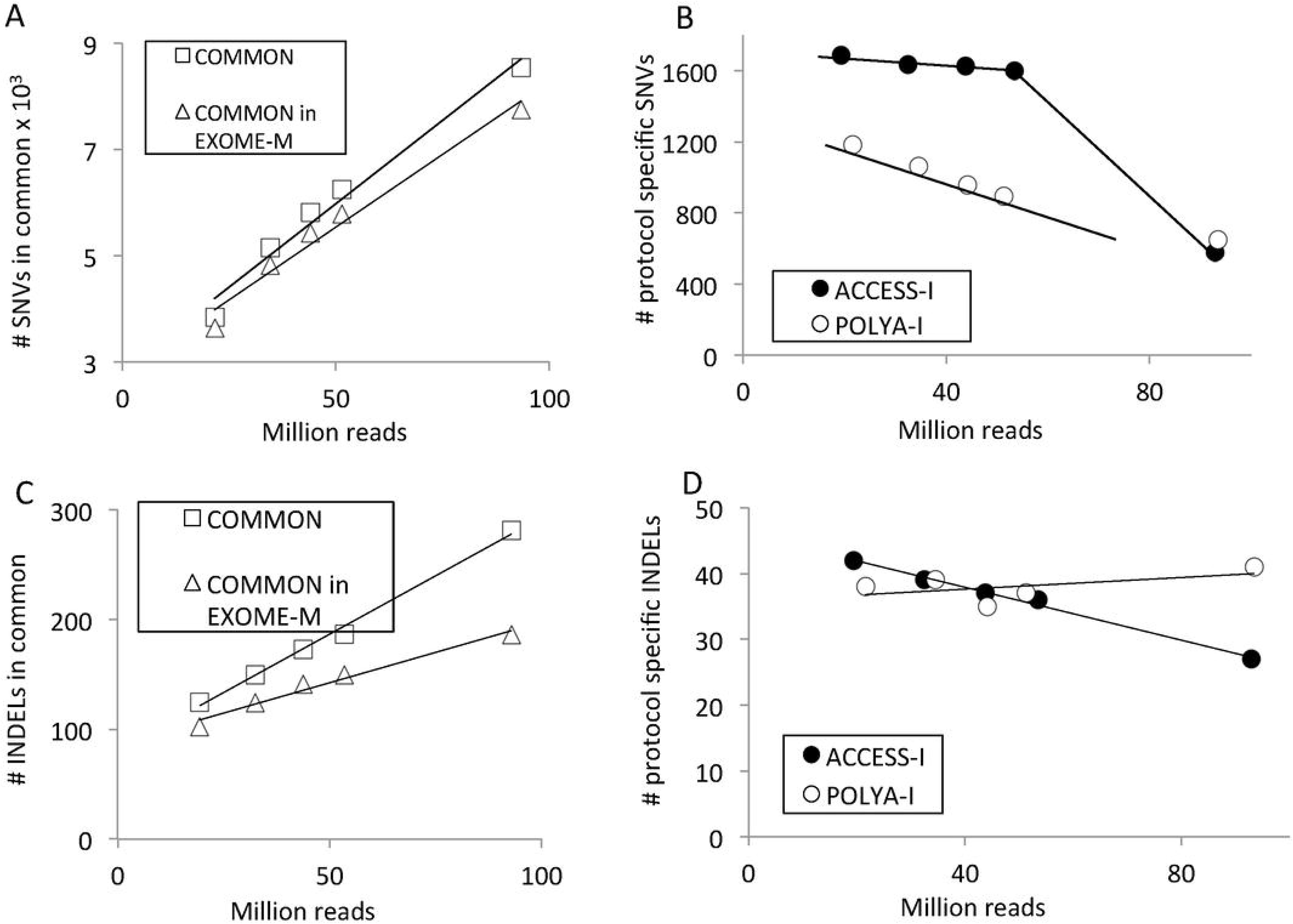
Coding SNVs/INDELs detected by ACCESS-I or POLYA-I protocol in function of increasing number of reads. A) Number of SNVs, in coding exons, detected in common between ACCESS-I and POLYA-I. B) Number of protocol specific coding SNVs also detected in MCF7 WES data. C) Number of INDELs, in coding exons, detected in common between ACCESS-I and POLYA-I. D) Number of protocol specific coding INDELs also detected in MCF7 WES data. ○ and ● Indicate, from left to right in each panel, s001, s012, s123, s1234,
SAB1234 samples.

To evaluate the coherence of protocol specific coding SNVs with WES data, we compared ACCESS-I and POLYA-I s001, s012, s123, s1234, sAB1234 results with respect to WES data of MCF7 cells (EXOME-M) previously published by Wang [2] (Supplementary Table 1).The EXOME-M dataset was analyzed as described in Fig. 1, and a total of 254668 SNVs were detected. ACCESS-I and POLYA-I SNVs in coding regions were intersected with EXOME-M SNVs. It is notable that the set of SNVs detected in common between ACCESS-I and POLYA-I were mostly included in the list of SNVs detected with WES in MCF7 cells (Fig. 5A Δ). However, when the subset of protocol specific SNVs, that were also included in MCF7 WES data, were analyzed, ACCESS protocol (Fig. 5B ●) came out to be able to detect more SNVs (at least > 400) than POLYA (Fig. 5B ○). The amount of ACCESS specific SNVs remained higher than the POLYA specific till 60 million sequenced reads (Fig. 5B), indicating that in a “standard” RNA-seq gene-level quantification experiment, that usually results in 30-40 million reads [19, 20], ACCESS might detect more coding SNVs with respect to POLYA.

The above analysis was also run for INDELs (Table 2, Fig. 5 C,D]. The number of INDELs in coding regions detected by the two protocols increased linearly with the increase of the number of sequenced reads (Table 2, Fig. 5C □). The subset of INDELs also present in WES data of MCF7 cells (EXOME-M, 218000 INDELs) increased linearly with the increment of sequencing reads, but with a flatter slope (Fig. 5C Δ) with respect to the one observed for SNVs (Fig. 5A Δ). Furthermore, when we analyzed the subset of protocol specific INDELs that were also included in WES data of MCF7 cells, ACCESS protocol (Fig. 5D ●) came out to be able to detect the same number of INDELs also detected by POLYA (Fig. 5D ○). The amount of ACCESS specific INDELs slightly decreased with the increase of the number of sequenced reads (Fig. 5D ●) as instead POLYA specific INDELs keep constant (Fig. 5D ○).

We have also intersected ACCESS-I and POLYA-I SNVs with the list of MCF7 SNVs annotated in COSMIC 77 database (Supplementary Figure 2). Also for the COSMIC 77 dataset the number of SNVs detected by ACCESS protocol was slightly higher than those detected by POLYA protocol.

### ACCESS and POLYA detect a similar number of fusion transcripts

Since RNA-seq is the preferred method for fusion transcripts detection [11], we also investigated the effect of ACCESS and POLYA protocols in this specific analysis, using the JAFFA method [17] to detect fusion transcripts. In the 5 sets of data samples generated from either POLYA-I or ACCESS-I datasets, the number of detectable fusion transcripts increased with respect to the number of mapped reads reaching a plateau at approximately 100 million reads (Table 3 and Fig. 6 ●○).

**Figure 6:**
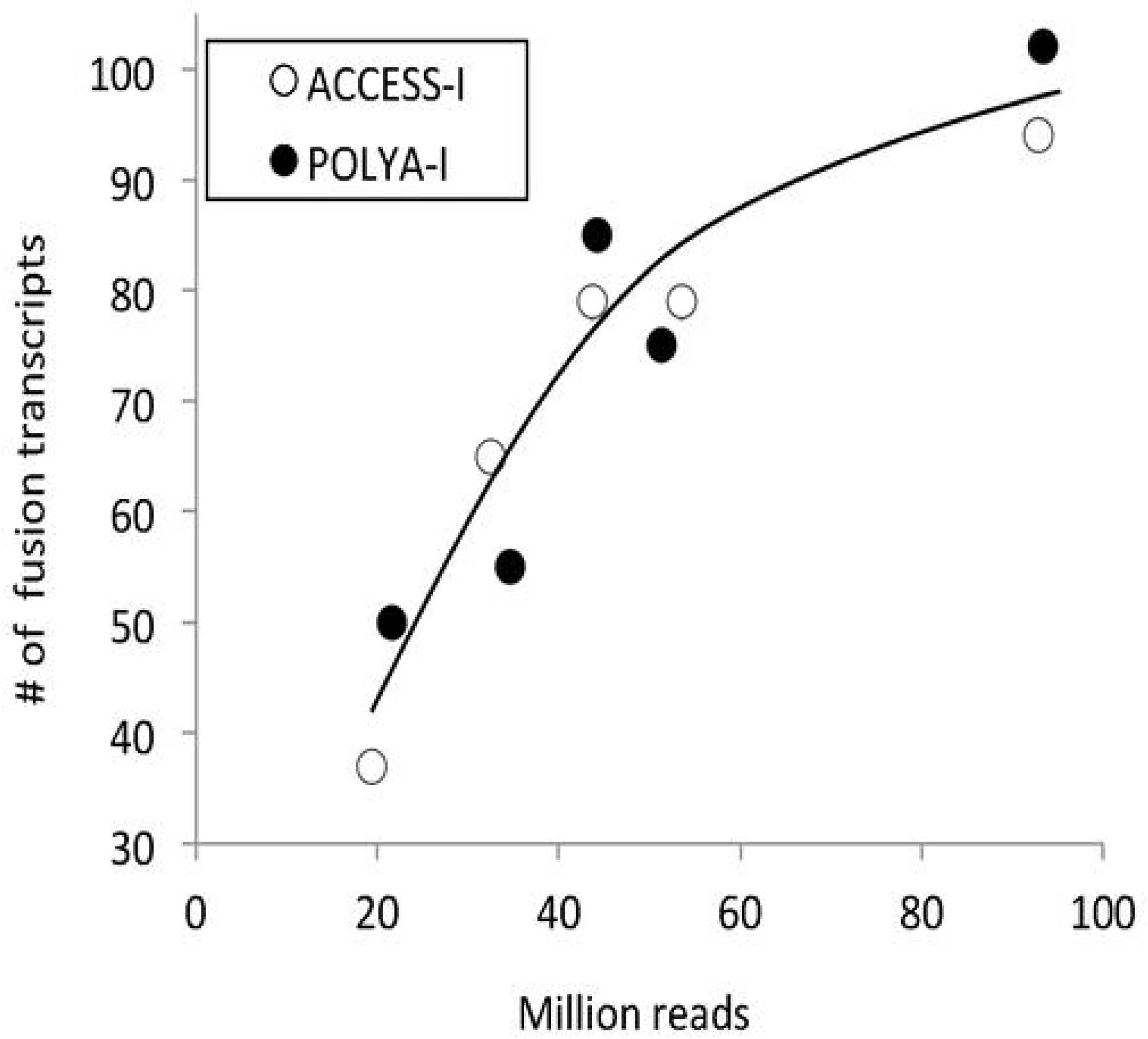
Fusion transcripts detected using ACCESS and POLYA protocols. Number of fusion transcripts detected is function of the number of sequenced reads.

Different Authors [6, 21–24] detected 3-41 fusion transcripts in the MCF7 cell line (Supplementary Table 2) and only 1-10 were in common. We compared the full list (52 fusion transcripts, Supplementary Table 3) to the list of those detected by us with Jaffa. As shown in Table 3 within the “already reported” fusion transcripts, both ACCESS and POLYA detected a similar number of fusion transcripts.

## Conclusions

We show here that the overlap between the SNV/INDELs detected by the two protocols under scrutiny was only partial. The reason of such partial overlap of the SNV/INDELs detected by the two protocols is intrinsic to the two protocols structure. ACCESS, which is based on an exons-specific capturing procedure, provided a better resolution for the SNV/INDELs located within coding exons, as instead since POLYA is based on full length polyA^+^ mRNA, it was particularly efficient in capturing SNV/INDELs associated to non-coding exons, i.e. 5’ and 3’ UTRs. The draw back of POLYA was the reduced sensitivity for coding SNVs in case the analysis is run on data collected for standard differential expression analysis, i.e. 30÷40 million reads. The needs of higher number of reads for POLYA is probably due to the large sampling pool characterizing polyA^+^ mRNAs, i.e. 221.9 Mb (calculated on the basis of the UCSC hg19 transcriptome), as instead ACCESS targets only coding exons, i.e. 37Mb (calculated on the basis of the ACCESS targeted regions).

Concerning fusion transcripts detection, the overall number of gene fusions detected by the two protocols is similar, but the detected fusion transcripts are only partially overlapping, indicating that the data structure due to different library preparation methods affects their detection. None of the two protocols has some particular advantage in the selective detection of previously know fusion events.

In conclusion, in case the RNA-seq analysis aims at detecting coding SNVs ACCESS might be preferred, as it requires less reads with respect to POLYA protocol. On the other hand, if the analysis aims also at identifying variants in the 5’and 3’ UTRs, POLYA protocol is definitively more suitable.

## Notes

Mariaflavia Di Renzo and Raffaele A. Calogero are both last authors.

## Declarations

### Acknowledgements

This work has been supported by: AIRC (Italian Association for Cancer Research) grants IG-17426 to AW and IG-17473 to MD) and Consiglio Nazionale delle Ricerche Flagship projects EPIGEN to RAC.

### Availability of data and materials

We thank Drs G. Schorth and S. Gross (Illumina, San Diego, CA) for providing MCF7 RNA-seq data generated with TruSeq RNA Sample Prep v2 and TruSeq RNA Access Library Prep Kits. These datasets are available upon request inquiring to. We also thanks Dr Y. Asmann (Mayo Clinic, Jacksonville, Florida) for providing MCF7 exome-seq data generated with Agilent’s SureCapture v2 kit, which can be requested to.

### Authors’ contributions

RAC and MD supervised the research. MA and MO designed the study. RP, FC and MB performed the bioinformatics analyses. RAC, MD and AW drafted the manuscript. All the authors read and approved the final manuscript.

## Figures legends

**Supplementary Figure 1:** Gene expression quantification by RSEM. A) Distribution of s001, s012, s123, s1234, sAB1234 log_2_ expression ratio between ACCESS-I and POLYA-I. 70% of the genes for the 5 datasets are included within +/- 0.5 range. B) Genes detected as consistently more expressed in ACCESS-I or POLYA-I using Rank Product analysis. Only the genes identified as more expressed in POLYA-I are expressed at least two folds more in POLYA-I than in ACCESS-I.

**Supplementary Figure 2:** A) Overlaps between SNVs detected in ACCESS-I and SNVs of MCF7 annotated in COSMIC 77 database, B) Overlaps between SNVs detected in POLYA-I and SNVs of MCF7 annotated in COSMIC 77 database

